# EvoMut: A Computational Framework for Engineering Oxidative Stability in Proteins

**DOI:** 10.64898/2026.03.19.712986

**Authors:** Seyed Shahriar Arab, Nathan E. Lewis

## Abstract

Amino acid oxidation is a major cause of protein instability and loss of function in therapeutic and industrial settings. Although methionine, cysteine, tyrosine, and tryptophan residues are widely recognized as oxidation-prone, only a subset of such residues are dominant functional hotspots, and not all are suitable targets for mutation. Identifying these vulnerable yet engineerable sites remains a major challenge. Here, we present EvoMut, a residue-level analytical framework for evaluating both oxidative vulnerability and mutation feasibility. EvoMut estimates oxidation risk by integrating structural features, local functional context, intrinsic chemical susceptibility, and evolutionary conservation. A central feature of the framework is the explicit separation of oxidation risk from mutation feasibility: candidate substitutions are evaluated only after high-risk residues are identified and ranked by evolutionary substitution patterns. Application of EvoMut to multiple proteins, and evaluation with experimental data, showed that oxidation-prone residues differ markedly in their engineering potential. EvoMut distinguishes residues that are both oxidation-sensitive and evolutionarily permissive from those that are chemically vulnerable but functionally constrained. By providing residue-level mechanistic insight, EvoMut offers a practical framework for the rational design of oxidation-resistant proteins. EvoMut is freely available as a web server at https://evomut.org.

**Significance Statement:** Strategies to improve oxidative stability in proteins often rely on chemical intuition or solvent accessibility alone, with limited consideration of functional and evolutionary constraints. EvoMut addresses this gap by explicitly separating oxidative vulnerability from mutation feasibility and integrating structural, chemical, functional, and evolutionary information within an interpretable framework. It helps explain why some oxidation-prone residues can be successfully engineered whereas others remain constrained; thus, supporting rational decision-making in oxidative stability engineering.

## Introduction

Oxidative modification is a pervasive phenomenon in protein science and represents a major source of structural instability, loss of enzymatic activity, and reduced functional lifetime in biological, industrial, and therapeutic contexts[1–4]. Reactive oxygen species (ROS) can damage proteins during production, formulation, storage, and *in vivo* circulation. Even subtle chemical modifications at specific residues may disrupt local interactions, alter folding behavior, promote aggregation, and ultimately compromise protein function.

At the molecular level, ROS preferentially target amino acids with high redox reactivity. Methionine and cysteine are among the most susceptible residues, and their oxidation can produce methionine sulfoxide or various cysteine derivatives that perturb tertiary structure and functional integrity[3]. Aromatic residues, such as tyrosine, may also undergo radical-mediated reactions that promote covalent cross-linking and aggregation[2,3]. Despite this chemical selectivity, experimental evidence consistently shows that oxidative damage is not uniformly distributed among oxidation-prone residues within a protein[5–7].

In many proteins, oxidative impairment is dominated by a limited number of critical sites. These residues are not necessarily the most solvent-exposed or intrinsically reactive in isolation; Instead, they occupy positions where structural context, functional coupling, and dynamic accessibility amplify the consequences of oxidative modification. Consequently, simple heuristics such as equating solvent accessibility with high oxidation risk or burial with protection often fail to explain experimental observations[8,9].

Protein engineering strategies aimed at improving oxidative stability have frequently focused on direct substitution of oxidation-prone residues with less reactive alternatives[10–12]. Although such approaches have been successful in specific cases, their outcomes remain highly variable and may lead to reduced catalytic activity, altered specificity, or compromised stability. This variability partly reflects the fact that protein stability and function emerge from coordinated interactions among neighboring residues, and modification of a single site can propagate indirect effects through local structural networks.

Two distinct factors therefore determine whether a residue represents a practical engineering target. The first is oxidative vulnerability, defined as the likelihood that a residue will undergo chemical modification in its native structural and functional environment. The second is mutation feasibility, which reflects whether substitution at that position is compatible with structural, functional, and evolutionary constraints.

A residue may be highly susceptible to oxidation yet remain functionally indispensable, severely restricting possible substitutions[5,12,13]. Conversely, some oxidation-prone residues may tolerate a broad range of substitutions with minimal functional consequences.

Here we introduce EvoMut, an integrative analytical framework that evaluates oxidative vulnerability and mutation feasibility as related but distinct residue-level properties. EvoMut estimates oxidation risk through a weighted integration of structural features, local functional context, intrinsic chemical susceptibility, and evolutionary conservation. For residues identified as high risk, the framework further evaluates mutation feasibility using evolutionary substitution patterns and structural contact relationships. By placing candidate substitutions within their broader structural and evolutionary context, EvoMut provides a systematic framework for identifying engineerable oxidation hotspots and guiding the rational design of oxidation-resistant proteins.

## Results

### EvoMut framework and design principles

In rational protein engineering, two fundamental questions frequently arise. First, which residues represent critical hotspots that strongly influence protein stability or function? Second, once such sites are identified, which substitutions can improve the desired property without disrupting structural integrity or biological activity? EvoMut was developed to address these questions in a systematic and sequential manner. The framework separates hotspot identification from substitution assessment and evaluates each stage using complementary sources of information. A schematic overview of the EvoMut workflow is shown in Figure 1.

**Figure 1.**
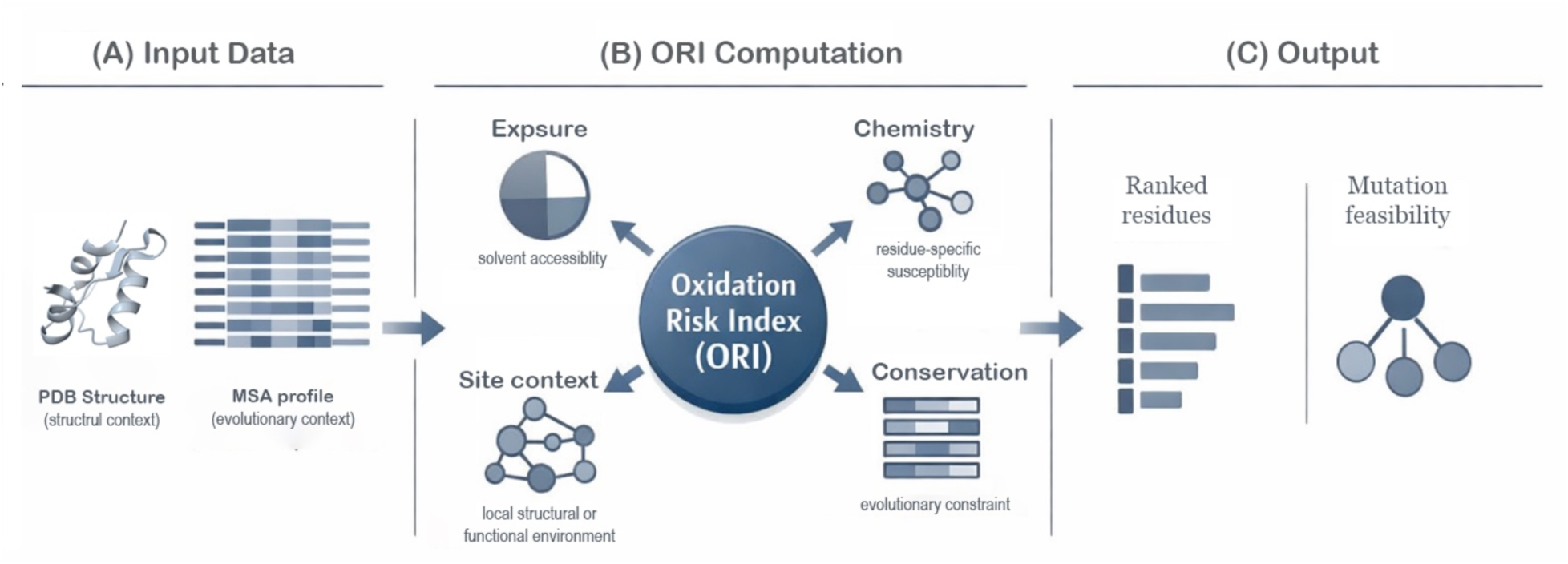
Schematic overview of the EvoMut framework. Protein structures obtained from the Protein Data Bank (PDB) are integrated with evolutionary sequence profiles derived from multiple sequence alignments (XSSP/HSSP). Residue-level oxidation susceptibility is estimated as a weighted combination of solvent accessibility, intrinsic chemical susceptibility, proximity to functional sites, and evolutionary conservation. Residues with elevated oxidation risk are subsequently evaluated for mutation feasibility using position-specific evolutionary substitution patterns, enabling prioritized identification of oxidation-sensitive yet evolutionarily tractable engineering targets.

### Identification of critical sites through multi-factor modeling of oxidation risk

In EvoMut, identification of oxidation hotspots is not based on a single structural or chemical feature. Instead, oxidation risk is modeled as a composite quantity derived from the weighted integration of several independent factors. These include solvent accessibility, intrinsic chemical susceptibility of the residue, proximity to annotated functional sites, and evolutionary conservation of the position. By combining these determinants, EvoMut avoids relying on simplified assumptions such as equating solvent accessibility with oxidation risk. Residues that are partially buried or located within structurally important regions may still emerge as high-risk positions when their functional context and chemical susceptibility are considered simultaneously.

Evolutionary information is incorporated at this stage with a relatively small weighting coefficient. This design ensures that strongly conserved residues are not overlooked when other factors suggest elevated susceptibility, while preventing conservation alone from dominating the risk assessment.

### Stage-specific use of evolutionary information for substitution assessment

After oxidation-sensitive residues are identified, EvoMut evaluates possible substitutions for each hotspot. At this stage, evolutionary information becomes the primary determinant of mutation feasibility. Position-specific substitution patterns derived from multiple sequence alignments are used to define the set of amino acids that are tolerated at each site. This approach restricts the substitution space to variants that are consistent with the evolutionary history of the protein and therefore more likely to preserve structural stability and function. Separating hotspot identification from substitution assessment prevents chemically reactive residues from being automatically interpreted as suitable mutation targets and ensures that engineering decisions remain consistent with evolutionary constraints.

### Role of neighboring residues and local structural context

Protein stability and function are rarely determined by a single residue in isolation. Instead, local networks of interacting residues collectively shape the structural environment of a site. EvoMut therefore evaluates residues that are spatially adjacent to each identified hotspot. Structural contact analysis is used to identify neighboring residues that may influence local stability or compensate for perturbations introduced by mutation. Evolutionary patterns of these neighboring positions are also examined to assess whether limited adjustments in the surrounding environment may improve the compatibility of a proposed substitution.

This context-aware analysis allows EvoMut to support both single-site mutation strategies and targeted multi-site adjustments when local structural interactions suggest that coordinated changes may be beneficial.

### Interpretability in support of engineering decisions

The central design objective of EvoMut is interpretability. Rather than producing opaque predictions, the framework reports residue-level results that can be traced back to contributing structural, chemical, and evolutionary factors. For each identified hotspot, the contribution of individual components to the calculated oxidation risk can be examined, and proposed substitutions are directly linked to observed evolutionary frequencies. This transparency allows EvoMut outputs to be interpreted as experimentally testable hypotheses that can guide rational protein engineering strategies

### 6.1 Global identification of oxidation-sensitive residues

Application of EvoMut to proteins with available structural and evolutionary information revealed that oxidative susceptibility is not uniformly distributed along protein sequences. Instead, a relatively small subset of residues consistently dominates oxidation-related functional vulnerability. Although several amino acids are chemically susceptible to oxidative modification, only a fraction of these residues emerge as dominant hotspots when structural context and evolutionary constraints are considered simultaneously. In many proteins, multiple oxidation-prone residues are present, yet only one or two positions account for most of the experimentally observed functional loss following oxidative stress.

Residues identified as high-risk sites typically exhibit a convergence of several contributing factors. These include intrinsic chemical susceptibility, structural exposure or functional proximity to catalytic regions, and evolutionary patterns that allow limited but plausible substitutions. When these determinants act together, certain positions display markedly elevated oxidation risk relative to other chemically-similar residues within the same protein.

This observation highlights the importance of integrative residue-level analysis. Methods based solely on solvent accessibility or residue chemistry often fail to explain why some oxidation-prone residues strongly affect protein activity while others remain functionally neutral. The following case studies illustrate how EvoMut distinguishes dominant oxidative hotspots from chemically reactive but functionally marginal residues and how evolutionary substitution patterns further clarify the feasibility of targeted substitutions.

### 6.2 Case Study I: Human α₁-Antitrypsin (A1AT)

Human α₁-antitrypsin (A1AT) was selected as a benchmark system because its oxidative inactivation has been extensively characterized experimentally. The protein contains eighteen residues that are chemically susceptible to oxidation, including nine methionines, six tyrosines, one cysteine, and two tryptophans. Despite this apparent abundance of oxidation-prone residues, experimental studies have shown that loss of inhibitory activity is primarily driven by oxidation at two specific positions, Met351 and Met358 within the reactive center loop[5,14,15].

Consistent with these observations, EvoMut analysis identified Met351 and Met358 as the dominant oxidation hotspot in A1AT. Both residues displayed substantially higher oxidation risk scores than the remaining methionines in the protein, forming a clearly separated upper tier in the risk distribution (Table 1). The complete residue-level ORI values are provided in Supplementary Table S2. Structural mapping further revealed that these residues are located within the reactive center loop, a region that plays a central role in the inhibitory mechanism of A1AT (Figure 2A).

**Figure 2.**
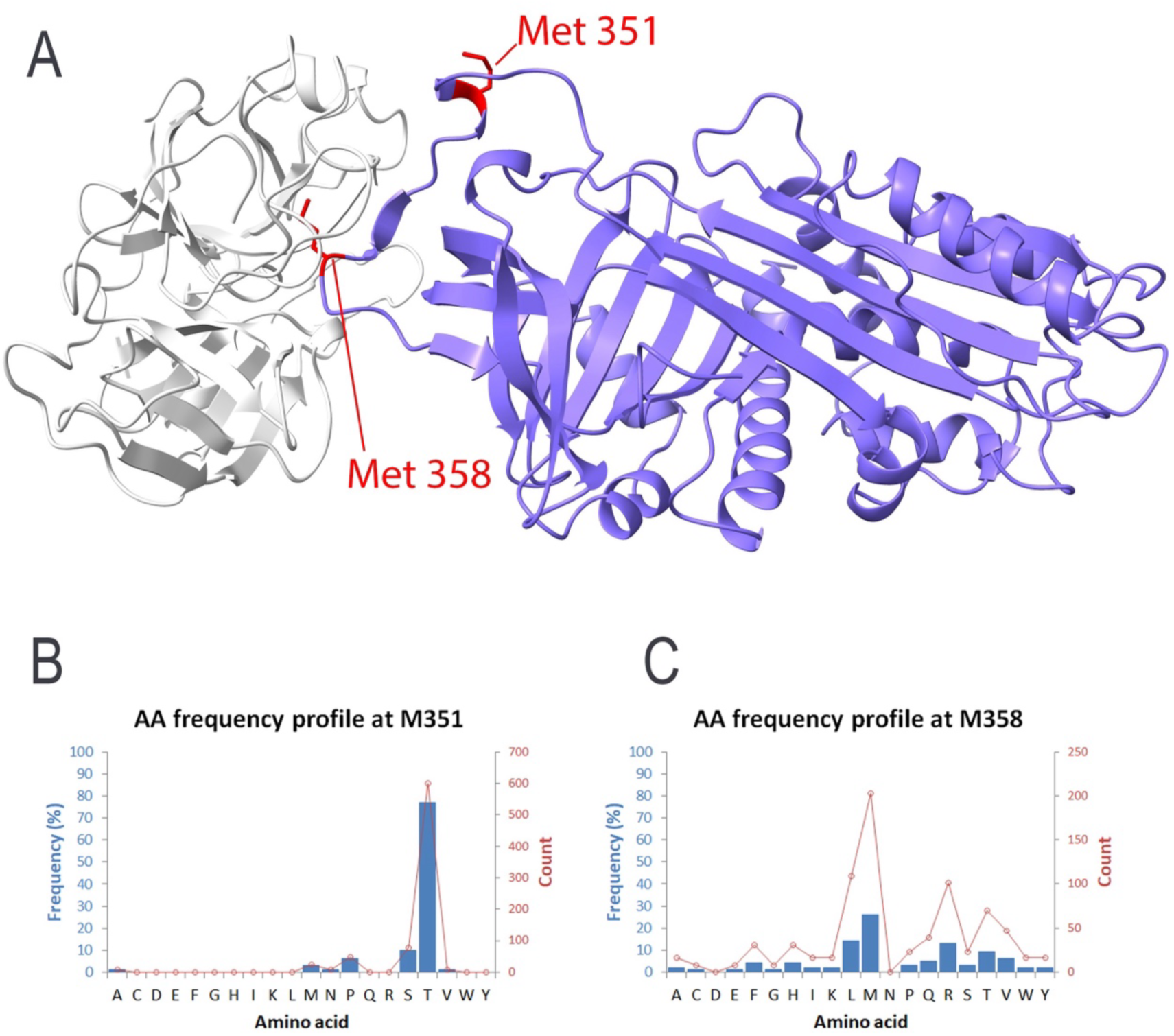
Structural and evolutionary analysis of Met351 and Met358 in A1AT (PDB: 1OPH). (A) Three-dimensional structure of A1AT highlighting Met351 and Met358 in red within the reactive center loop. (B) Evolutionary residue- frequency distribution at position 351 showing strong preference for threonine (∼77%) with methionine observed at low frequency (∼3%), indicating restricted substitution tolerance. (C) Evolutionary residue-frequency distribution at position 358 showing broader substitution tolerance, with methionine (∼26%) and several alternative residues present.

**Table 1.** Residue-level oxidation risk profile for human α₁-antitrypsin. Oxidation risk index (ORI) values are reported for all oxidation-prone residues in the protein sequence. Met358 and Met351 form a distinct upper tier in the risk distribution, reflecting the combined influence of structural context, solvent accessibility, and proximity to the reactive center loop.

| Position | WT | WT freq. | ORI |
| --- | --- | --- | --- |
| **358** | **M** | **26** | **0.95** |
| **351** | **M** | **3** | **0.89** |
| 226 | M | 21 | 0.73 |
| 374 | M | 3 | 0.50 |
| 242 | M | 30 | 0.46 |
| 385 | M | 29 | 0.46 |
| 63 | M | 69 | 0.40 |
| 221 | M | 97 | 0.35 |
| 220 | M | 98 | 0.35 |
| 238 | W | 27 | 0.34 |
| 160 | Y | 73 | 0.23 |
| 194 | W | 99 | 0.21 |
| 187 | Y | 82 | 0.21 |
| 297 | Y | 94 | 0.20 |
| 38 | Y | 89 | 0.19 |
| 244 | Y | 98 | 0.18 |
| 138 | Y | 99 | 0.18 |

Evolutionary analysis provided additional insight into the functional constraints at these two positions. The residue-frequency distribution derived from sequence alignments indicates that position 351 is strongly biased toward threonine (∼77%), with methionine appearing only at low frequency (∼3%) (Figure 2B). This restricted substitution pattern suggests that structural or functional requirements limit the range of tolerated residues at this site. In contrast, position 358 displays a broader evolutionary substitution spectrum. Methionine remains present in approximately 26% of sequences, while several alternative residues including leucine, valine, isoleucine, threonine, and glutamine occur at appreciable frequencies (Figure 2C). This pattern indicates a comparatively higher tolerance for amino-acid replacement at this position.

Together, these results establish a clear residue-level hierarchy within A1AT. EvoMut identifies Met351 and Met358 as the dominant oxidation-sensitive hotspots while simultaneously revealing distinct differences in their evolutionary substitution tolerance. This distinction highlights an important aspect of oxidative stability engineering: residues that are similarly vulnerable to oxidation may differ substantially in their suitability as mutation targets.

### 6.3 Case Study II: Industrial α-Amylases

#### 6.3.1 Alkaline α-Amylase from *Alkalimonas amylolytica*

The alkaline α-amylase (AmyA) from Alkalimonas amylolytica is an industrial enzyme widely used in detergent and textile applications, where exposure to oxidative agents such as hydrogen peroxide can substantially reduce catalytic performance. In a structure-guided engineering study by Yang et al., several methionine residues located near the catalytic core were individually substituted with leucine to improve oxidative stability[8]. Because the experimental study reports residue numbers based on sequence annotation, whereas our analysis uses PDB-based structural numbering, the corresponding positions are mapped as follows: Met247→349, Met214→316, Met229→331, Met317→421, and Met145→235. For clarity, structural numbering is used throughout the following analysis.

All single Met→Leu variants displayed improved oxidative stability relative to the wild-type enzyme. Among these mutations, substitution at position 349 (Met247 in the experimental numbering) produced the most pronounced improvement, retaining approximately 72% of catalytic activity after exposure to 500 mM H₂O₂.

Independent EvoMut analysis identified Met349 as the highest-risk oxidation site among all methionine residues in AmyA (ORI = 0.88). Evolutionary profiling further revealed a restricted substitution space at this position, dominated by hydrophobic residues, with leucine representing the most frequently observed alternative. Consistent with this evolutionary permissibility, EvoMut ranked leucine as the optimal substitution candidate. Structural analysis shows that both residues are located near the catalytic region, yet they differ markedly in their evolutionary profiles (Figure 3A). Methionine is highly conserved at position 316 (∼96%), indicating strong evolutionary constraint, whereas position 349 displays lower methionine conservation and a broader substitution spectrum (Figure 3B–C). This contrast suggests that mutation at position 316 may be structurally less tolerated despite its moderate solvent accessibility.

**Figure 3.**
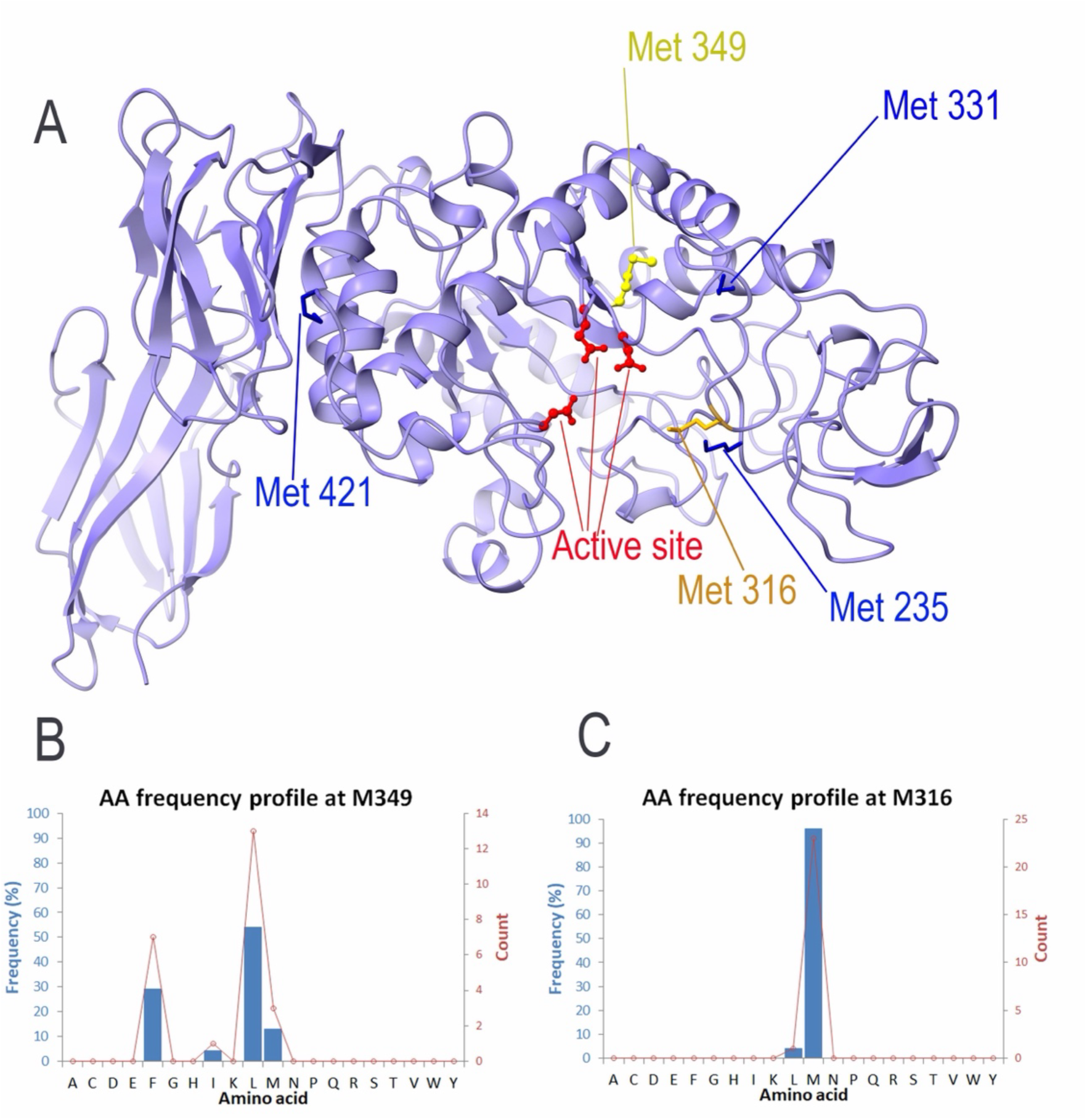
Structural and evolutionary analysis of methionine residues in alkaline α-amylase (PDB: 3BC9). (A) Three- dimensional structure showing active-site residues in red. Met349 (yellow) and Met316 (orange) are positioned in proximity to the catalytic region, whereas Met235, Met331, and Met421 are shown in blue. (B) Evolutionary residue-frequency distribution at position 349, dominated by leucine (54%) and phenylalanine (29%), with methionine present at lower frequency (13%), indicating moderate substitution tolerance. (C) Evolutionary residue- frequency distribution at position 316, where methionine is strongly conserved (96%) and leucine is observed at low frequency (4%), reflecting strong evolutionary constraint at this position.

Taken together, these observations illustrate two important principles. First, oxidation vulnerability cannot be inferred from solvent accessibility alone; residues that appear buried may still represent dominant functional hotspots when their structural context and catalytic proximity are considered. Second, evolutionary conservation provides a critical constraint on mutation feasibility. By integrating these factors, EvoMut distinguishes functionally important oxidation hotspots from residues that are structurally constrained but less influential in oxidative inactivation.

The strong agreement between experimental structure-guided engineering and EvoMut’s residue-level prioritization highlights Met349 as the dominant oxidative vulnerability in AmyA and demonstrates the utility of integrative structural and evolutionary analysis for guiding oxidation- resistance engineering.

#### 6.3.2 α-Amylase AmyC from *Thermotoga maritima*

The intracellular α-amylase AmyC from Thermotoga maritima was analyzed to evaluate site-specific variation in oxidation susceptibility and substitution tolerance at methionine residues that have been experimentally investigated. In a previous mutagenesis study, Ozturk et al. examined four positions (Met43, Met44, Met55, and Met62) and reported that substitution of Met55 with alanine (M55A) produced the most pronounced improvement in oxidative stability and enzymatic performance among the tested variants[9].

Consistent with these experimental observations, EvoMut analysis ranked Met55 as the most oxidation-sensitive residue among the examined positions. The calculated oxidation risk index (ORI) was highest for Met55 (0.53), followed by Met43 (0.51), whereas Met62 and Met44 showed intermediate and nearly comparable values (0.48). These results indicate that position 55 represents the dominant oxidation hotspot within the experimentally investigated set.

Structural analysis further shows that Met55 is embedded within a predominantly hydrophobic local environment formed by neighboring residues (Figure 4B). This structural context suggests that oxidative modification at this position may disrupt local packing interactions, thereby affecting enzyme stability.

**Figure 4.**
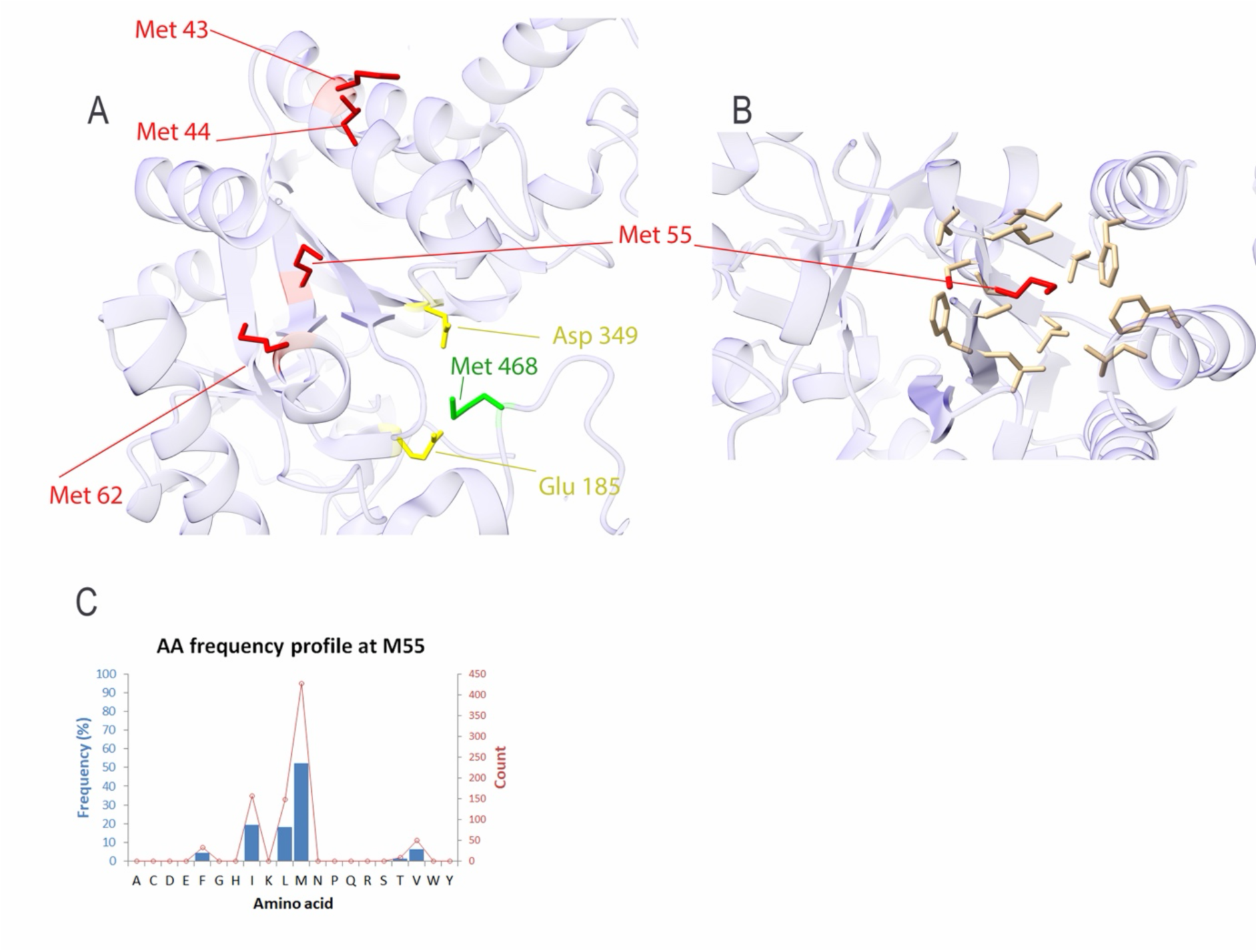
Structural and evolutionary characterization of methionine residues in α-amylase AmyC (PDB: 2B5D). (A) Three-dimensional structure of α-amylase AmyC showing experimentally investigated methionine residues Met43, Met44, Met55, and Met62 in red. The catalytic residues Asp349 and Glu185 are shown in yellow, and Met468 is highlighted in green. (B) Local structural environment of Met55 (red) with neighboring residues shown in yellow, illustrating a predominantly hydrophobic local environment. (C) Evolutionary residue-frequency distribution at position 55, showing partial conservation of methionine (52%) and enrichment of hydrophobic residues including isoleucine (19%), leucine (18%), and valine (6%), consistent with structural constraints at this position.

Evolutionary residue-frequency analysis provides additional insight into substitution tolerance at this site. The full residue-frequency distribution derived from the HSSP alignment is provided in Supplementary Table S3. Position 55 shows partial conservation of methionine (∼52%), while several hydrophobic residues, including isoleucine (∼19%), leucine (∼18%), and valine (∼6%), are also observed in the evolutionary profile (Figure 4C). The prevalence of hydrophobic residues indicates that maintenance of a nonpolar side chain is structurally favored at this position.

In contrast, Met43 and Met44 display broader evolutionary substitution patterns that include polar and charged residues, suggesting that these sites are subject to weaker structural constraints. Met62 exhibits a more restricted pattern but still shows lower predicted oxidation risk than Met55.

Substitution risk profiling across the amino-acid alphabet further indicates that isoleucine is predicted to provide the greatest reduction in oxidation susceptibility at position 55 while maintaining structural compatibility with the local environment. This prediction differs from the experimentally tested alanine substitution but highlights an alternative design strategy in which hydrophobic replacements preserve the packing interactions surrounding the hotspot.

Taken together, these results identify Met55 as the primary oxidation-sensitive site among the experimentally examined residues in α-amylase AmyC and illustrate how EvoMut can refine mutation strategies by integrating oxidation risk with evolutionary substitution constraints.

### 6.4 Case Study III: Versatile Peroxidase (VP)

To further evaluate the ability of EvoMut to interpret experimentally characterized oxidation-sensitive systems, we analyzed the versatile peroxidase (VP) engineering dataset reported by Sáez- Jiménez et al.[12]. In that study, several methionine residues located near the heme cofactor and catalytic environment were investigated through site-directed mutagenesis to determine their contribution to oxidative inactivation.

EvoMut analysis of the VP structure revealed multiple residues with elevated susceptibility to oxidative modification. Notably, the four methionine residues that were experimentally targeted in the original study (M152, M247, M262, and M265) were all identified among the oxidation-prone positions in the EvoMut analysis, enabling a direct comparison between computational predictions and experimental observations.

Within this subset, EvoMut assigned oxidation risk index (ORI) values of 0.69 for M247, 0.66 for M152, 0.63 for M265, and 0.62 for M262. These values fall within a relatively narrow range, indicating that oxidative susceptibility in VP is distributed across several methionine residues rather than being dominated by a single critical hotspot.

This pattern is consistent with the experimental findings, where individual substitutions produced only moderate improvements in oxidative stability. No single mutation was sufficient to substantially enhance resistance to oxidative inactivation, suggesting that multiple residues contribute collectively to the observed sensitivity.

From an engineering perspective, this result illustrates an important design principle. When predicted oxidation risk is distributed across several residues with comparable ORI values, mutation of a single site is unlikely to produce a large stabilization effect. Instead, effective stabilization strategies may require coordinated modification of multiple oxidation-prone residues within the catalytic environment.

Taken together, these findings demonstrate that EvoMut can capture both types of oxidative vulnerability patterns: systems dominated by a single critical hotspot, as observed in A1AT and AmyA, and systems in which susceptibility is distributed across several residues, as observed in versatile peroxidase. This capability is particularly valuable for guiding rational engineering strategies in proteins where oxidative sensitivity arises from multiple contributing sites.

## Discussion

Oxidative modification represents a major source of functional instability in many proteins, yet the relationship between chemical susceptibility and practical engineering targets is often complex. The analysis presented here demonstrates that residues with high intrinsic oxidation susceptibility do not necessarily represent the most effective mutation targets. Instead, the functional impact of oxidative modification depends strongly on structural context and evolutionary constraints.

Across the systems examined in this study, EvoMut consistently identified a limited subset of residues that dominate oxidative vulnerability. In A1AT, oxidation risk was concentrated at Met351 and Met358, both located within the reactive center loop (Figure 2). These residues displayed markedly elevated oxidation risk compared with other methionines in the protein (Table 1), consistent with experimental studies demonstrating that oxidation at either site disrupts inhibitory activity. At the same time, evolutionary analysis revealed substantial differences in substitution tolerance between the two sites. Position 351 exhibits a highly restricted evolutionary profile, whereas position 358 shows a broader substitution spectrum. This contrast illustrates that residues with similar oxidative sensitivity may differ significantly in their mutational flexibility.

A similar pattern was observed in the industrial α-amylase from Alkalimonas amylolytica. EvoMut ranked Met349 as the dominant oxidation-sensitive site, in agreement with experimental mutagenesis studies in which substitution at this position produced the largest improvement in oxidative stability. Notably, this residue is structurally buried, whereas other methionines with greater solvent accessibility exhibited lower predicted oxidation risk. This observation highlights a key limitation of heuristic approaches that rely primarily on solvent accessibility when identifying oxidation-sensitive residues. Structural context and functional proximity can outweigh solvent accessibility in determining the functional consequences of oxidative modification.

The analysis of the thermophilic α-amylase AmyC further illustrates how evolutionary information can refine mutation strategies. Experimental mutagenesis identified Met55 as the most influential oxidation-sensitive position, and substitution with alanine improved oxidative stability. However, the evolutionary profile of this site shows that hydrophobic residues with larger side chains, such as isoleucine, leucine, and valine, are frequently observed at this position. This pattern suggests that substitutions preserving both hydrophobic character and side-chain volume may better maintain local packing interactions than smaller residues such as alanine (Figure 4). Integrating evolutionary substitution patterns with structural context, provides additional guidance for selecting alternative mutation strategies beyond those initially explored experimentally.

In contrast to the previous examples, analysis of versatile peroxidase revealed a distributed pattern of oxidative vulnerability. Several methionine residues exhibited comparable oxidation risk values, and experimental mutagenesis confirmed that individual substitutions produced only moderate improvements in oxidative stability. This pattern suggests that oxidative sensitivity in such systems arises from the combined contribution of multiple residues rather than a single dominant hotspot. In these cases, effective stabilization strategies may require coordinated modification of several oxidation-prone residues within the catalytic environment.

Taken together, these results highlight two general principles that govern oxidation-driven protein instability and guide rational engineering strategies. First, oxidation-sensitive hotspots emerge from the combined influence of chemical susceptibility, structural context, and functional coupling rather than from residue chemistry alone. Second, evolutionary substitution patterns provide a powerful constraint for identifying mutations that are more likely to preserve structural integrity and activity.

By integrating these complementary sources of information, EvoMut enables residue-level interpretation of oxidative vulnerability while simultaneously restricting the mutational search space to biologically plausible substitutions. This approach allows protein engineers to prioritize a small number of experimentally testable variants, reducing the need for extensive screening while maintaining mechanistic interpretability in protein engineering workflows.

## Conclusion

Oxidative modification remains a major challenge in the stability and long-term functionality of many proteins used in biomedical and industrial applications. The analyses presented here demonstrate that oxidative vulnerability cannot be reliably predicted from residue chemistry or solvent accessibility alone. Instead, susceptibility emerges from the combined influence of structural context, functional proximity, and evolutionary constraints.

EvoMut provides an interpretable residue-level framework that separates the identification of oxidation-sensitive hotspots from the evaluation of mutation feasibility. By integrating structural features with evolutionary substitution patterns, the framework enables prioritization of mutation strategies that are both chemically meaningful and biologically plausible.

Application of EvoMut to experimentally-characterized systems illustrates how this integrative approach can distinguish dominant oxidative hotspots from structurally constrained residues and clarify when single-site or multi-site engineering strategies may be required. Together, these capabilities support design of oxidation-resistant proteins while reducing the experimental search space in protein engineering projects.

## Methods

### Implementation

EvoMut is implemented as a web-accessible analytical framework written in Python and designed with a modular architecture to enable flexible residue-level analysis. The implementation separates data ingestion, structural analysis, evolutionary annotation, risk scoring, and mutation assessment into independent computational modules, allowing each stage of the workflow to be evaluated or extended without affecting the overall framework.

Structural features are extracted directly from the input protein structure using established bioinformatics libraries for parsing coordinate files and analyzing residue-level properties. Solvent accessibility, secondary structure, and inter-residue contacts are computed from the three-dimensional coordinates. Evolutionary information is incorporated through residue-specific profiles derived from multiple sequence alignments.

All intermediate calculations are handled in memory during runtime to ensure session isolation and reproducibility. The framework is deployed as an interactive web application that integrates numerical analysis with synchronized sequence-level and structure-level visualization, allowing users to explore oxidation risk and mutation feasibility in an iterative manner.

### Input Data

#### Protein structure

EvoMut requires a three-dimensional protein structure obtained from the Protein Data Bank (PDB). Users may provide a PDB identifier or upload a local coordinate file. For each structure, EvoMut automatically detects available chains and extracts the amino-acid sequence directly from atomic coordinate records, ensuring consistency between structural and sequence representations.

Residue-level structural features are derived from the structure, including relative solvent accessibility, secondary structure assignment, and spatial proximity between residues. These features are subsequently used to evaluate residue exposure, identify structurally relevant regions, and determine neighboring residues within a user-defined distance threshold.

### Evolutionary profile

Evolutionary information is incorporated through residue-specific sequence profiles derived from multiple sequence alignments. EvoMut supports HSSP-formatted profiles, which can be retrieved from the XSSP server (https://www3.cmbi.umcn.nl/xssp/)[16].

These profiles provide position-specific amino-acid frequencies and conservation metrics that reflect evolutionary constraints at each residue. Within EvoMut, evolutionary data serve two complementary purposes: they contribute modestly to oxidation risk estimation by penalizing highly conserved positions, and they guide the identification of plausible amino-acid substitutions based on observed evolutionary variability.

Optional annotations, such as known functional residues or sequence features, may also be incorporated when available in UniProt annotations to refine site-context evaluation, although they are not required for analysis.

### Oxidation Risk Assessment

For each residue, EvoMut estimates oxidation susceptibility by integrating structural, chemical, and evolutionary determinants into a unified scoring framework. The oxidation risk index (ORI) is defined as a weighted combination of multiple normalized factors:

ORI = *w_E_*⋅E + *w_C_*⋅C + *w_K_*⋅(1-K) + *w_S_*⋅S

where

E represents relative solvent accessibility (RSA),

C represents intrinsic chemical susceptibility of the amino acid,

K represents the evolutionary frequency of the residue in the multiple sequence alignment,

S represents the functional-site context score, reflecting proximity to annotated catalytic or binding residues.

The coefficients w_E_, w_C_, w_K_, and w_S_ define the relative contribution of each factor. All variables are normalized to the range between 0 and 1, resulting in a continuous per-residue ORI value between 0 (low oxidation susceptibility) and 1 (high susceptibility).

Default weighting parameters are provided in Supplementary Table S1. These values can be interactively adjusted through the EvoMut interface to emphasize structural or evolutionary contributions depending on the characteristics of the protein system under investigation.

### Mutation Feasibility Assessment

Residues identified as high-risk oxidation hotspots are subjected to a second-stage analysis focused on mutation feasibility. At this stage, evolutionary information plays a central role. Position-specific substitution patterns derived from multiple sequence alignments are used to determine which amino-acid replacements are evolutionarily tolerated at each site. Candidate substitutions are therefore restricted to residues that occur within the observed evolutionary substitution spectrum.

Substitutions are subsequently evaluated based on their reduced chemical susceptibility relative to the native residue while maintaining compatibility with evolutionary constraints. This staged approach ensures that chemically favorable substitutions are also consistent with the structural and functional constraints reflected in the evolutionary record.

### Analysis of Local Structural Context

Protein stability and function often arise from networks of interactions among neighboring residues rather than from the behavior of a single residue in isolation. EvoMut therefore evaluates residues that are spatially adjacent to each identified hotspot. Structural contact analysis is used to identify neighboring residues within a defined spatial cutoff. Evolutionary patterns of these surrounding residues are examined to assess whether limited adjustments within the local environment may improve compatibility with the proposed substitution.

This analysis supports context-aware mutation strategies in which small adjustments within local interaction networks may mitigate destabilizing effects or preserve catalytic functionality.

### Output and Visualization

EvoMut produces residue-level outputs that are fully traceable and interpretable. These include oxidation risk scores for all residues in the protein, identification of high-risk oxidation hotspots, evolutionarily supported substitution candidates for each hotspot, and contextual information about neighboring residues and local structural contacts.

Results are presented through an interactive interface that integrates tabular summaries, sequence-level annotation, and three-dimensional structural visualization. These synchronized views allow users to examine oxidation risk and mutation feasibility within their structural and evolutionary context. All output tables can be exported for downstream analysis or experimental planning. Screenshots illustrating the EvoMut workflow and analysis interface are provided in Supplementary Figures S1–S3.

### Availability

The EvoMut framework is available as a web server at https://evomut.org. The datasets supporting this study are publicly available in Zenodo at https://doi.org/10.5281/zenodo.19039897. Additional methodological details and supporting material are provided in the Supplementary Information.

### Scope of Applicability

EvoMut is designed for protein engineering scenarios in which residue-level decisions are required to guide targeted mutational interventions. The framework is particularly suited for systems in which oxidative inactivation is driven by a limited number of critical residues.

By restricting candidate substitutions to those supported by evolutionary evidence and structural context, EvoMut reduces the mutational search space and prioritizes experimentally testable hypotheses when extensive screening is impractical.

## Acknowledgements

This work was supported by NIGMS (R35 GM119850), NHLBI (R41 HL164260), the Novo Nordisk Foundation (NNF20SA0066621) and the Georgia Research Alliance.

## Notes

https://evomut.org/

https://doi.org/10.5281/zenodo.19039897

### Competing Interest Statement

The authors have declared no competing interest.

