## Supplementary Information for "EvoMut: A Computational Framework for Engineering Oxidative Stability in Proteins"

### Supplementary Methods

#### Sup. Method 1. Definition of the Oxidation Risk Index (ORI)

The oxidation risk index (ORI) used in EvoMut quantifies residue-level susceptibility to oxidative modification as a weighted combination of multiple independent determinants. The ORI score is defined as:

$$\text{ORI} = w_E \cdot E + w_C \cdot C + w_K \cdot (1 - K) + w_S \cdot S$$

where:

**E (Exposure)**
Relative solvent accessibility (RSA) derived from structural analysis.

**C (Chemical liability)**
Residue-type–specific intrinsic susceptibility to oxidative modification. Amino acids such as methionine, cysteine, tyrosine, and tryptophan receive higher baseline susceptibility scores due to their known chemical reactivity toward reactive oxygen species.

**S (Site context)**
Proximity of the residue to annotated functional or binding sites derived from UniProt annotations or structural interface analysis.

**K (Conservation)**
Normalized evolutionary conservation score derived from multiple sequence alignment entropy.

The coefficients $w_E$, $w_C$, $w_K$, and $w_S$ define the relative contribution of each factor. All coefficients are scaled to the range **[0,1]** prior to integration. Default weighting coefficients used throughout this study are summarized in Supplementary Table S1.

These values were selected to reflect the dominant contributions of intrinsic chemical susceptibility and structural context while allowing evolutionary conservation to modulate, but not dominate, oxidation risk estimation.

#### Sup. Method 2. Evolutionary substitution analysis and mutation feasibility

Evolutionary substitution profiles were derived from XSSP/HSSP multiple sequence alignments, which provide residue-specific amino acid frequencies across homologous protein sequences.

For each residue position, amino acid frequencies were extracted and normalized to represent the observed evolutionary tolerance of substitutions at that site.

Mutation feasibility was evaluated only after identification of residues with elevated oxidation risk and was treated as a separate analytical stage. Candidate substitutions were ranked based on:

- Observed evolutionary frequency at the target position
- Structural compatibility inferred from the local residue environment (e.g., surface-exposed vs buried residues)
- Absence of known pathogenic annotations when available (e.g., ClinVar annotations for human proteins)

This staged design ensures that oxidation susceptibility and mutation feasibility are evaluated independently, preventing chemically reactive residues from being automatically interpreted as suitable mutation targets.

#### Sup. Method 3. Identification of functional and interface residues

Functional context contributes to oxidation risk estimation through the site context component (S) of the ORI score.

For multimeric protein structures, EvoMut automatically identifies residues that participate in inter-chain contacts. Residues located within a defined distance threshold from atoms belonging to another chain are classified as interface residues and contribute to the functional-site context score.

For single-chain proteins, EvoMut can optionally incorporate manually specified functional residues such as catalytic or binding residues when available. These annotations may be derived from UniProt functional features or experimental literature.

This flexible design allows EvoMut to integrate both automatically detected structural interfaces and manually annotated functional sites to refine residue-level context evaluation.

### Supplementary Figures

#### Sup. Fig. 1: Step 1 (Input data)

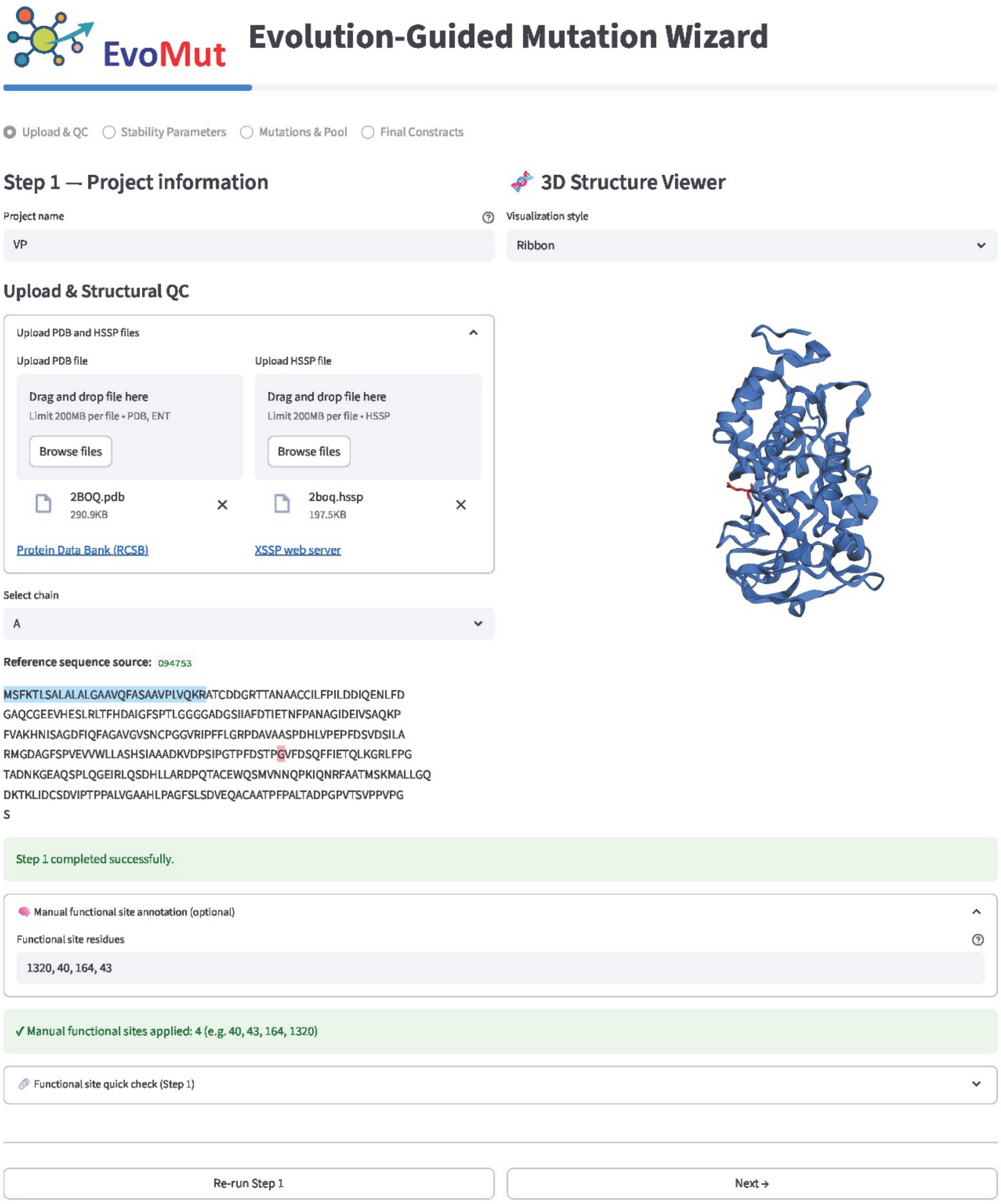

*Supplementary Figure S1 EvoMut web interface – Step 1: Input configuration. Users provide the PDB structure, HSSP alignment, and chain identifier used for the analysis. Optional parameters allow specification of reference sequence and functional-site residues when required.*

### Sup. Fig. 2: Step 2 (Parameters setting)

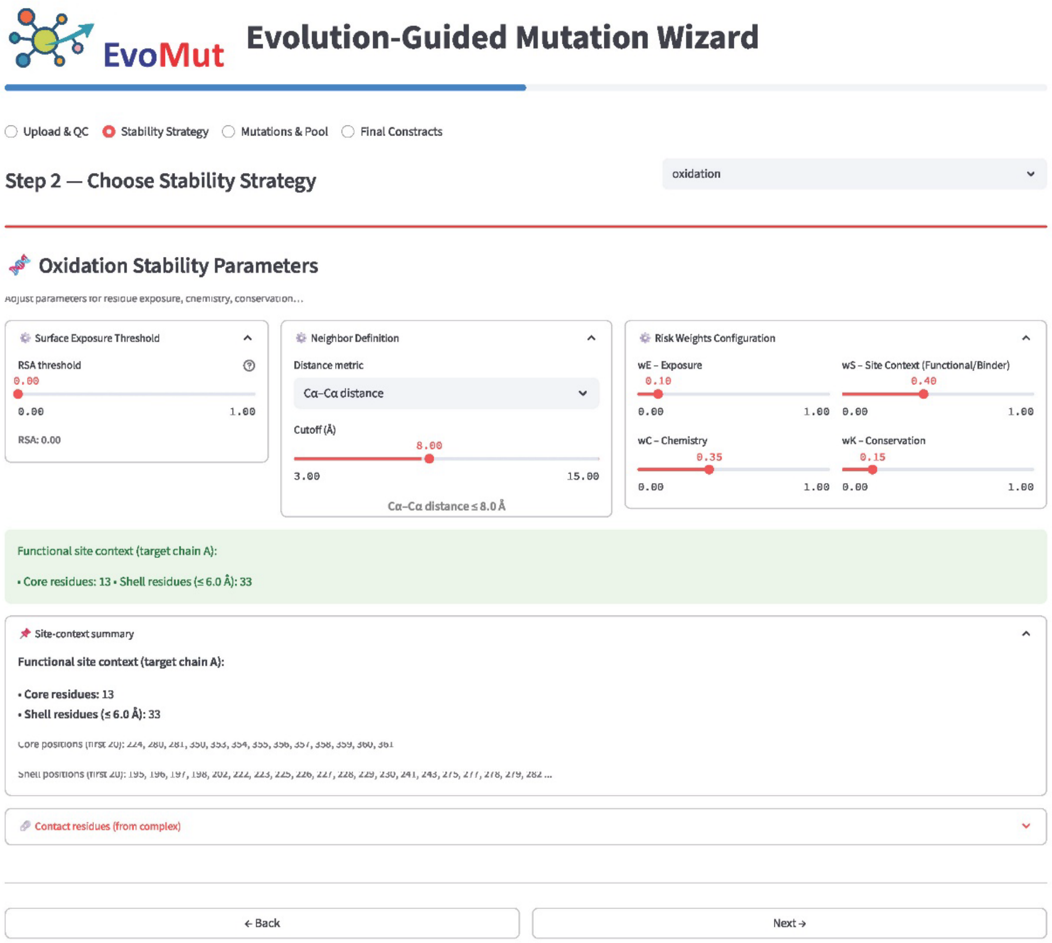

*Supplementary Figure S2 EvoMut web interface – Step 2: Structural analysis. The program evaluates residue-level structural properties including solvent accessibility, spatial neighborhood, and proximity to functional or interface residues.*

### Sup. Fig. 3: Step 3 (Analysis data and building constructions)

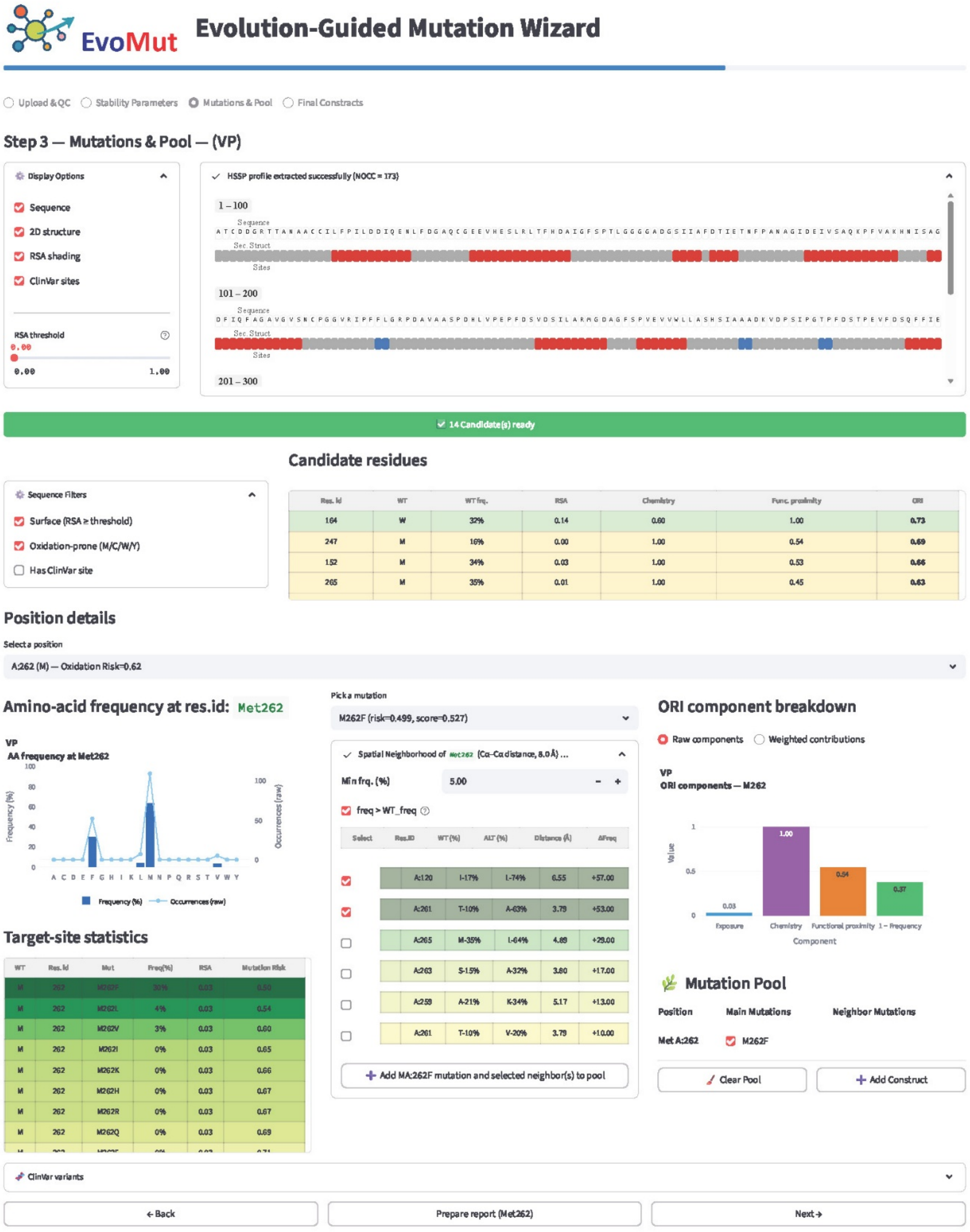

*Supplementary Figure S3 EvoMut web interface – Step 3: Oxidation risk and mutation analysis. The EvoMut framework calculates oxidation risk scores and identifies candidate substitutions based on evolutionary substitution profiles.*

### Supplementary Tables

#### Supplementary Table S1. Default weighting coefficients

*Supplementary Table S1 Default weighting coefficients used in the ORI scoring function.*

| Parameter | Description | Default value |
|---|---|---|
| $w_E$ | Solvent exposure weight | 0.10 |
| $w_C$ | Chemical susceptibility weight | 0.35 |
| $w_S$ | Functional site context weight | 0.40 |
| $w_K$ | Evolutionary conservation weight | 0.15 |

#### Supplementary Table S2. Oxidation-prone candidates' info

*Supplementary Table S2 Complete oxidation risk index (ORI) values for oxidation-prone residues in human $\alpha_1$-antitrypsin.*

| Pos. | WT | WT frq. | RSA | Chemistry | Func. proximity | ORI |
|---|---|---|---|---|---|---|
| 358 | M | 26 | 0.88 | 1.0 | 1.00 | 0.95 |
| 351 | M | 3 | 0.56 | 1.0 | 0.83 | 0.89 |
| 226 | M | 21 | 0.13 | 1.0 | 0.63 | 0.73 |
| 374 | M | 3 | 0.01 | 1.0 | 0.00 | 0.50 |
| 242 | M | 30 | 0.04 | 1.0 | 0.00 | 0.46 |
| 385 | M | 29 | 0.01 | 1.0 | 0.00 | 0.46 |
| 63 | M | 69 | 0.00 | 1.0 | 0.00 | 0.40 |
| 221 | M | 97 | 0.00 | 1.0 | 0.00 | 0.35 |
| 220 | M | 98 | 0.00 | 1.0 | 0.00 | 0.35 |
| 238 | W | 27 | 0.22 | 0.6 | 0.00 | 0.34 |
| 160 | Y | 73 | 0.19 | 0.5 | 0.00 | 0.23 |
| 194 | W | 99 | 0.01 | 0.6 | 0.00 | 0.21 |
| 187 | Y | 82 | 0.06 | 0.5 | 0.00 | 0.21 |
| 297 | Y | 94 | 0.15 | 0.5 | 0.00 | 0.20 |
| 38 | Y | 89 | 0.01 | 0.5 | 0.00 | 0.19 |
| 244 | Y | 98 | 0.00 | 0.5 | 0.00 | 0.18 |
| 138 | Y | 99 | 0.01 | 0.5 | 0.00 | 0.18 |

#### Supplementary Table S3. Evolutionary residue-frequency distribution

*Supplementary Table S3 Evolutionary residue-frequency distribution for Met55 in α-amylase AmyC derived from the HSSP alignment.*

| AA | Frequency_percent | Count_raw | Mutation_Risk |
|---|---|---|---|
| A | 0 | 0 | 0.84 |
| C | 0 | 0 | 0.97 |
| D | 0 | 0 | 0.79 |
| E | 0 | 0 | 0.71 |
| F | 4 | 33 | 0.60 |
| G | 0 | 0 | 0.92 |
| H | 0 | 0 | 0.67 |
| I | 19 | 156 | 0.46 |
| K | 0 | 0 | 0.66 |
| L | 18 | 148 | 0.46 |
| M | 52 | 427 | |
| N | 0 | 0 | 0.78 |
| P | 0 | 0 | 0.78 |
| Q | 0 | 0 | 0.70 |
| R | 0 | 0 | 0.68 |
| S | 0 | 0 | 0.84 |
| T | 1 | 8 | 0.72 |
| V | 6 | 49 | 0.57 |
| W | 0 | 0 | 0.94 |
| Y | 0 | 0 | 0.83 |

### Supplementary Data Availability

Complete EvoMut analysis outputs, including structural input files, HSSP alignments, configuration files, and detailed analysis results, are available in the Zenodo repository:

https://doi.org/10.5281/zenodo.19039897

Each case study described in the manuscript is organized in a separate directory containing the input data and EvoMut output files used in the analysis.
